# Underreported in-water behaviours of the loggerhead sea turtle: *Foraging on sea cucumbers*

**DOI:** 10.1101/2023.03.25.534220

**Authors:** Kostas Papafitsoros

## Abstract

In this article, we report on four observations of loggerhead sea turtles (*Caretta caretta*) foraging on sea cucumbers (*Holothuria spp*.), on Zakynthos Island, Greece. Direct observations of this behaviour are underreported in the literature. Foraging on such prey was in general a challenging process as the turtles used their flippers and their beak to cut the sea cucumbers albeit without success. They ended up shallowing the sea cucumbers most likely in one intact piece. Even though this behaviour is rare in this site, two out of four observations involved the same male sea turtle, indicating potentially specific dietary preferences for this individual.

## Introduction

The present article continues the series of short articles *Underreported in-water behaviours of the loggerhead sea turtle* which was initiated in (Papafitsoros, 2022b). The purpose of this series is to describe in-water behaviours of loggerhead sea turtles (*Caretta caretta*) that have been rarely reported in the literature or have not been reported at all. All these reports are the result of the author’s long-term (13 years) in-water study and photo-identification of the loggerhead sea turtles of Laganas Bay, Zakynthos, Greece. Laganas Bay hosts one of the most important reproductive grounds for the Mediterranean loggerheads (Casale *et al*., 2018) with about 300 females (Margaritoulis *et al*., 2022) and 100 males (Schofield *et al*., 2017) breeding annually. In-water observations, citizen science and photo-identification have also revealed the existence of a small year-round resident population (approximately 40 individuals) consisting mainly by adult males and juveniles (Papafitsoros *et al*., 2021; Papafitsoros *et al*., 2023). These individuals can be regularly seen foraging on sponges (*Chondrilla nucula*), molluscs (Papafitsoros and Schofield, 2016; Schofield *et al*., 2022), as well as discarded fishermen bycatch at a small port in Laganas Bay, (Agios Sostis port; author’s personal observations).

Here we report on four observations focusing on three individual turtles foraging on sea cucumbers (*Holothuria spp*.). Even though sea cucumbers have been found in the digestive tract of deceased turtles in multiple occasions, direct observations of loggerhead sea turtles preying on sea cucumbers have only been reported once in the literature (Rogers *et al*., 2020). We believe that the present article provides some interesting insights on these foraging events particularly with respect to their rarity, the difficulty of foraging on sea cucumbers as well as potential individual foraging specialisation on such prey.

## Methods

The study site is situated in Laganas Bay, Zakynthos, Greece (37^*◦*^ 43’N, 20^*◦*^ 52’E), Figure 1, which hosts an important breeding site for the Mediterranean loggerhead sea turtles (Casale *et al*., 2018). long-term in-water surveys combined with photoidentification (based on the sea turtles’ unique facial scales) have also shown that Laganas Bay is also a foraging habitat for around 40 resident turtles, mostly males and juveniles (Schofield *et al*., 2020; Papafitsoros *et al*., 2021). The present report is a result of these long-term in-water surveys. In summary, surveys involved an observer swimming close to the shore (max 7 metres depth), who, upon encountering a turtle, would take photographs for photo-identification purposes and to record any interesting behaviours. We refer e.g. to (Schofield *et al*., 2020) for more details on the methods and general context.

**Figure 1.**
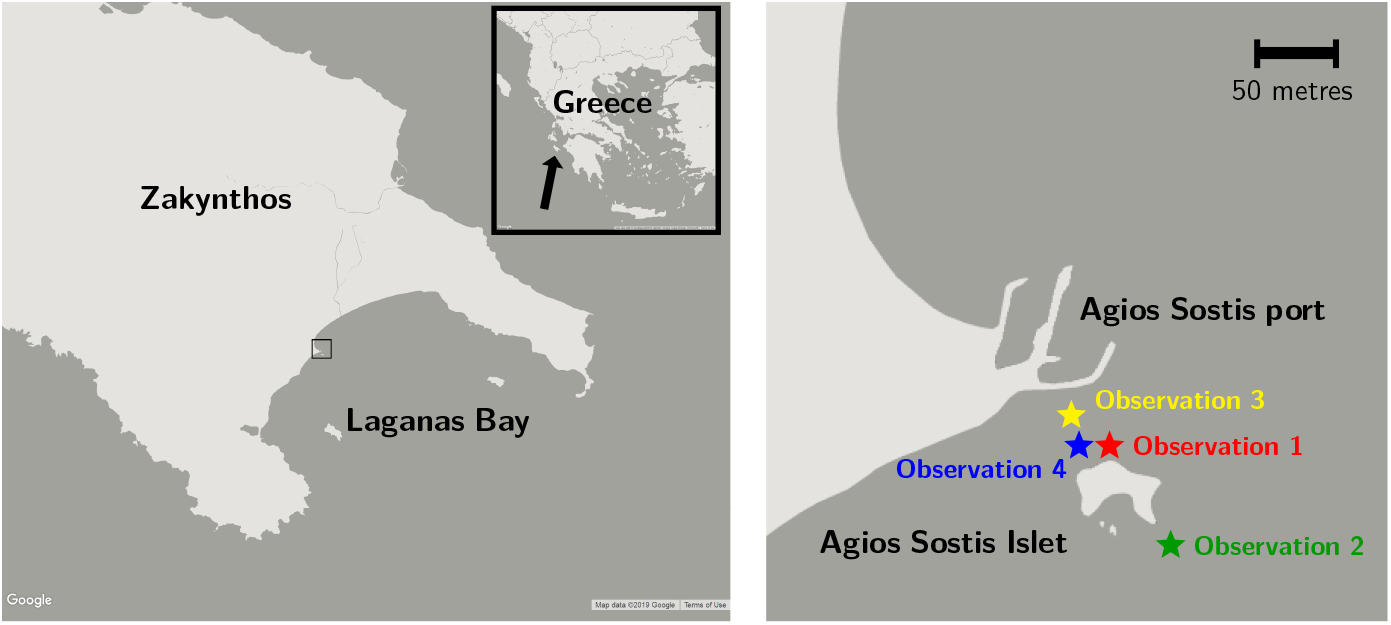
Left: Map of Greece and Laganas Bay, Zakynthos. Right: Map which corresponds to the area enclosed by the small square on the left map, showing the approximate locations of the four observations near Agios Sostis area at the northwest part of Laganas Bay.

## Results

All four observations took place in Agios Sostis area, around Agios Sostis Islet at the northwest part of Laganas Bay, Zakynthos, see Figure 1 for the approximate locations. The reef around Agios Sostis Islet is a well-known foraging ground in which resident male and juvenile turtles of Zakynthos prey on the sponge *Chondrilla nucula* which is abundant on this reef, (Papafitsoros and Schofield, 2016; Schofield *et al*., 2022). All the three turtles in the observations below have been such long-term residents and they have been logged in a existing photo-identification database consisting of more than 1500 individuals from this site, (Schofield *et al*., 2020; Papafitsoros *et al*., 2021).

### Observation 1

This observation took place on 26 August 2017 at the north side of Agios Sostis Islet foraging reef and involved the loggerhead turtle with ID name “t217” (possibly juvenile; estimated straight carapace length: 60-70cm). This individual was a regular occupier of the reef from July 2015 until October 2018. It was consistently observed foraging on sponges during that period (June-October) and this constitutes the only time it was observed foraging on a sea cucumber. The turtle initially observed around 07:36 and at 07:37:56 it spotted a sea cucumber (*Holothuria spp*., possibly *Holothuria poli*) at an approximate depth of 2 metres, see Figure 2. The turtle began the foraging attempt and tried to use its front flippers and its beak in order to cut it into smaller pieces, albeit without succeeding so. In the process it dropped the sea cucumber twice. It managed to shallow the sea cucumber, most likely in one intact piece in just under two minutes (around 07:39:43), and proceeded in foraging on sponges thereafter.

**Figure 2.**
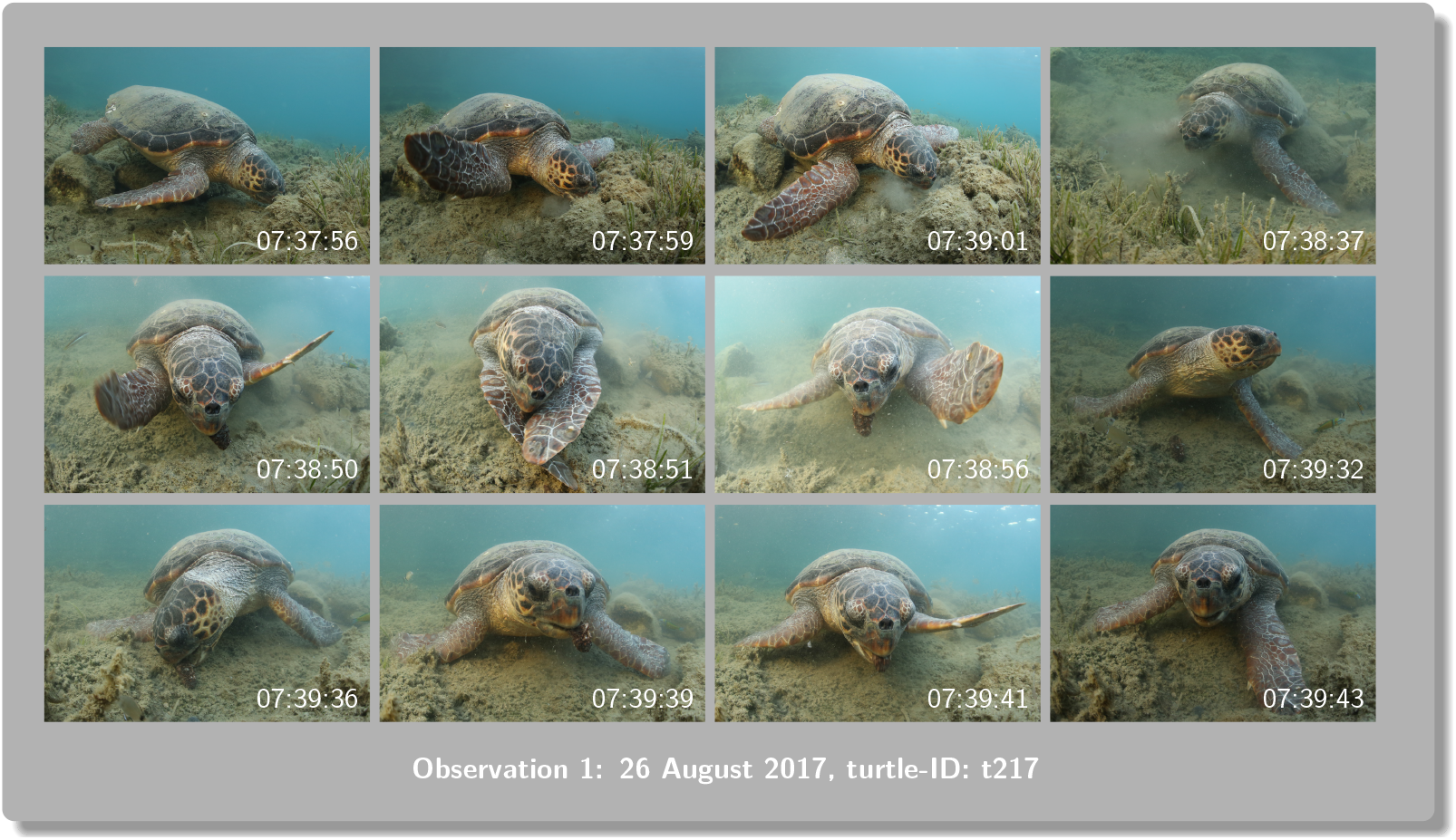
Successive photographs of Observation 1. The times on the bottom left represent the local times that each photograph was taken, as it was internally recorded by the camera.

### Observation 2

This observation took place on 15 June 2019 approximately 50m south of the Agios Sostis Islet foraging reef, and involved the loggerhead turtle with ID name “G18-04” (male, possibly adult; estimated straight carapace length: 70-80cm). The turtle has been a resident of Zakynthos from 2018 until at least the summer of 2022 and while it has been seen around the reef, it has not been observed foraging on sponges like the previous turtle. The turtle was first encountered swimming east of the reef at 07:51 and it spotted and started foraging on a sea cucumber (again *Holothuria spp*., possibly *Holothuria poli*) at 08:11:06 at an approximate depth of 8 metres, see Figure 3. Upon an initial closer inspection, the sea cucumber was observed being cut in half but it could not be determined whether this was due to turtle’s bites or whether it had been found in this state by the turtle. The turtle dropped the sea cucumber at least once and it could be seen alternatingly protruding from its mouth and being totally inside, indicating a difficulty from the turtle’s side to bite it and cut it. The turtle managed to shallow the sea cucumber at 08:15:21, just after 4 minutes, most likely again in one intact piece. However, it could not be excluded that it foraged on more than one sea cucumbers during this time period, since our observation was intermittent due to the large depth.

**Figure 3.**
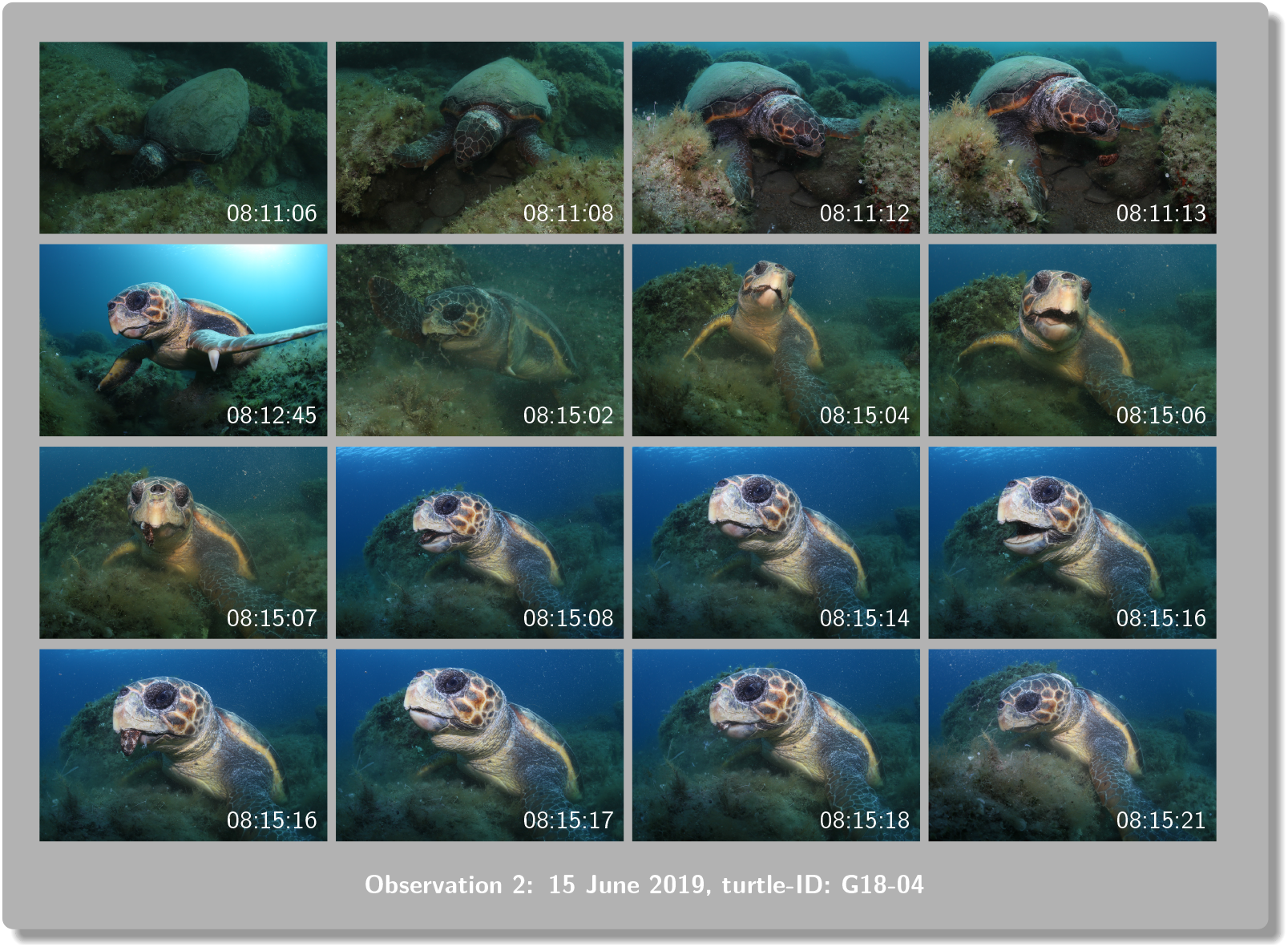
Successive photographs of Observation 2. The times on the bottom left represent the local times that each photograph was taken, as it was internally recorded by the camera. 06:51:28 06:51:28 06:51:29 06:51:34

### Observation 3

The third observation took place on 16 June 2019, that is, one day after Observation 3. It involved the same male loggerhead with ID name “G18-04”. It was observed at 07:51, on the north side of Agios Sostis Islet foraging reef, near the same location of Observation 1. The turtle was initially spotted holding already a sea cucumber in its mouth (*Holothuria spp*.), at an approximate depth of 2 metres, see Figure 4. At the same time a second loggerhead approached it (ID name “t323”). The second turtle was also a long-term occupant of the reef, foraging consistently on sponges, from 2016 until 2020, see also (Papafitsoros, 2022a). The male turtle “G18-04” began making circular avoiding movements, eventually releasing the sea cucumber. No physical contact occurred between the two individuals. The two turtles separated after 2 minutes, after which “G18-04” was briefly followed, swimming away from the spot. We did not check whether the second turtle attempted to forage on the sea cucumber.

**Figure 4.**
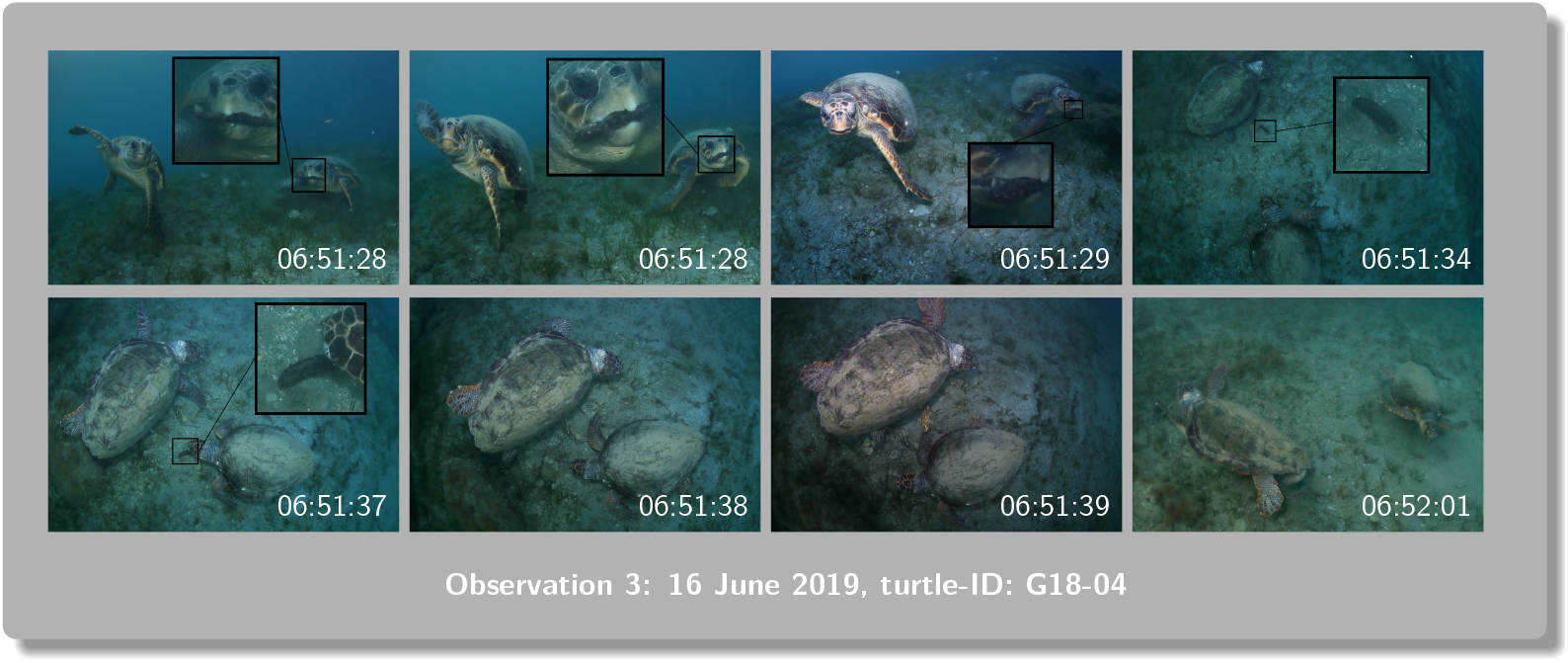
Successive photographs of Observation 3. The times on the bottom left represent the local times that each photograph was taken, as it was internally recorded by the camera. 18:30:01 18:30:02 18:30:05 18:30:07

### Observation 4

The fourth observation took place on 8 September 2020, again very close to the location of Observations 1 and 3. It involved the loggerhead sea turtle with ID name “t441” (possibly juvenile; estimated straight carapace length: 55-65cm) who has been a occupier of the reef from July 2019 until at least 2022. This turtle has also been consistently foraging on sponges throughout that period and similarly to the turtle of Observation 1, this was the only time it was observed foraging on a sea cucumber. It was spotted around 18:30 while it was attempting to forage on a sea cucumber (*Holothuria spp*., possibly *Holothuria poli*), at an approximate depth of 2 metres, see Figure 5. The turtle was also being observed by a couple of snorkelers but their presence did not seem to affect its behaviour, see also (Papafitsoros, 2015). Similarly to the turtle in Observation 1, it extensively used its front flippers attempting to cut the sea cucumber, again without success, and it dropped it at least six times during that process. The turtle managed to shallow the sea cucumber, again most likely in one intact piece, after about 3.5 minutes, and proceeded in foraging on sponges thereafter.

**Figure 5.**
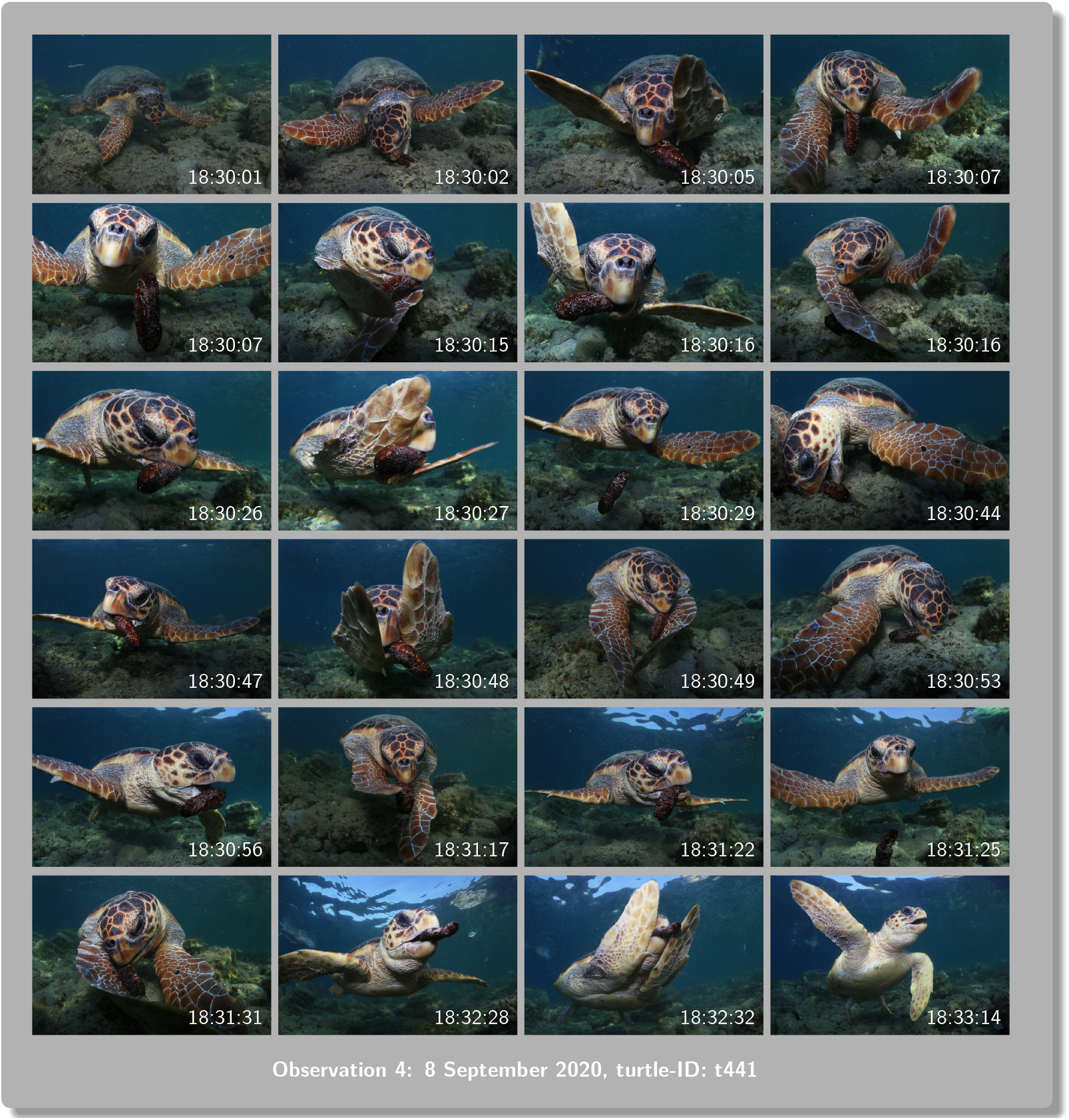
Successive photographs of Observation 4. The times on the bottom left represent the local times that each photograph was taken, as it was internally recorded by the camera.

Detailed photographs of all the four sea cucumbers can be found in Figure 6.

**Figure 6.**
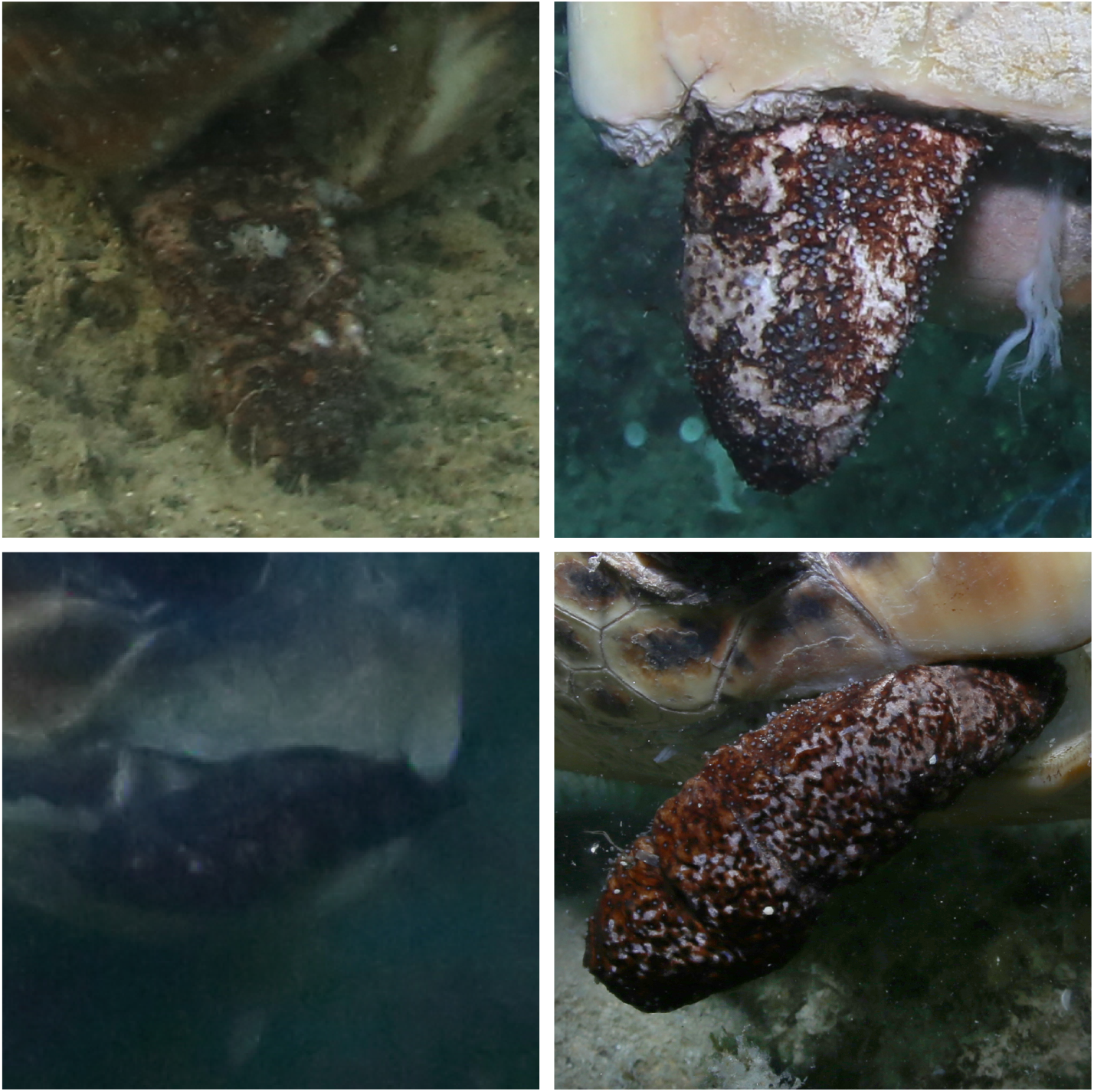
Detailed photographs of the four sea cucumbers (*Holothuria spp*., possibly *Holothuria poli*).

## Discussion

Sea cucumbers have been found in the digestive tract of Mediterranean loggerhead sea turtles during necropsies but the frequency of their occurrence is relatively low. For instance, Casale *et al*., 2008 analysed the gut contents of 79 turtles from the central Mediterranean sea, and only one (occurrence 1.3%) had consumed animals from the class *Holothuroidea*. Lazar *et al*., 2002 performed necropsies on four loggerheads from northern Adriatic sea and sea cucumbers represented 3% of the total dry weight of their prey. Similarly, Mariani *et al*., 2023, examined the gastrointestinal contents of 150 loggerhead sea turtles stranded in the central Adriatic Sea and central Tyrrhenian sea. The frequency of occurrence of animals of the Phylum Echinodermata (to which sea cucumbers belong) was founds to be 8.9% and 3.3% for turtles from the two areas respectively. Similar frequencies for *Holothuroidea* (3.3%) were found by Karaa *et al*., 2018, after performing necropsies on 132 loggerheads from the Gulf of Gabes, Tunisia. Further reports can be also found in Palmer *et al*., 2021, where also a photograph of a possibly intact sea cucumber recovered from a turtle’s ingestive tract is provided.

This relatively low frequency of consumption of sea cucumbers reported in the above studies, reflects the rarity of observations of this consumption in our study site. Indeed, 13 years of in-water surveys in Zakynthos Island, have resulted to only four such observations even though loggerheads can be regularly seen foraging on other prey (sponges). One simple explanation could be that sea cucumbers are in low abundance in our study site. However, we also observed that turtles exhibited difficulties in consuming such a prey, which aligns with the report by Rogers *et al*., 2020. Turtles failed to cut the sea cucumbers into pieces, and they most likely ended up shallowing it intact, a process that can take more than 4 minutes. We observed that this difficulty can lead to unsuccessful attempts, especially due to intraspecific interactions (Observation 3) which are very common in our study site (Schofield *et al*., 2022). In contrast, Mulochau *et al*., 2021 reported the case of a juvenile green sea turtle (*Chelonia mydas*) from Reunion island, which was observed foraging on two specimen of the sea cucumber *Synapta maculata* and it successfully used its beak and flippers to cut it into pieces. This was perhaps facilitated by the typical serrated jaw that green sea turtles have.

Finally, the fact that two of these rare observations involved the same loggerhead “G18-04”, which also avoided foraging on the abundant sponges that the other turtles in the area forage on, could be indicative of potentially specific dietary preferences of this individual. It has been argued that individual loggerheads can exhibit longterm foraging specialization (Vander Zanden *et al*., 2010; Hall *et al*., 2015; Pajuelo *et al*., 2016), a specialization that has also been observed in other sea turtles species, e.g. in green turtles (Vander Zanden *et al*., 2013). For instance, as an interesting example, the authors in Vicente and Carballeira, 1991, reported the gut contents of six hawksbill sea turtles (*Eretmochelys imbricata*) from Puerto Rico, with one adult only having ingested the sea cucumber *Holothuria cubana*, where the stomachs of the other turtles contained only sponges. It has been argued that long-term foraging specialization among others could reduce intraspecific competition (Vander Zanden *et al*., 2010). This could explain the dietary preferences of the individual “G18-04”, since turtles foraging on sponges on that reef are regularly involved in aggressive interactions (Schofield *et al*., 2022). For instance, “G18-04” was observed in that reef during the previous year (2018) when it was attacked by the individual “t217” (Observation 1). Notably the latter individual had been the most aggressive turtle of the reef for these two years (2018-2019), (Schofield *et al*., 2022). Specialising on a less abundant, and difficult to locate prey, like sea cucumbers, could lead to reduced engagement in intense energy demanding intraspecific interactions over foraging resources. Such specialisation indeed seems to occur in the resident population of Zakynthos with some individuals consistently foraging on sponges, whereas others focus on mining for molluscs in nearby submerged sandbanks (author’s personal observations).

## References

Casale, P., Abbate, G., Freggi, D., Conte, N., Oliverio, M., & Argano, R. (2008). Foraging ecology of loggerhead sea turtles Caretta caretta in the central Mediterranean Sea: Evidence for a relaxed life history model. Marine Ecology Progress Series 372, 265–276. https://doi.org/10.3354/meps07702.

Casale, P., Broderick, A.C., Camiñas, J.A., Cardona, L., Carreras, C., Demetropoulos, A., Fuller, W.J., Godley, B.J., Hochscheid, S., Kaska, Y., Lazar, B., Margaritoulis, D., Panagopoulou, A., Rees, A.F., Tomás, J., &Türkozan, O. (2018). Mediterranean sea turtles: current knowledge and priorities for conservation and research. Endangered Species Research 36, 229–267. https://doi.org/10.3354/esr00901.

Hall, A.G., Avens, L., McNeill, J.B., Wallace, B., & Goshe, L.R (2015). Inferring long-term foraging trends of individual juvenile loggerhead sea turtles using stable isotopes. Marine Ecology Progress Series 537, 265–276. https://doi.org/10.3354/meps11452.

Karaa, S., Imed, J., & Bradai, M. (2018). Diet of the Loggerhead Sea Turtles in th e Gulf of Gabes (Southern Tunisia, Central Mediterranean Sea). Poster presented at the 6th Mediterranean Conference on Marine Turtles, Poreç, Croatia. https://www.researchgate.net/profile/Karaa-Sami/publication/341965535_Diet_of_the_loggerhead_sea_turtles_in_the_Gulf_of_Gabes_Southern_Tunisia_Central_Mediterranean_Sea/links/5edb474b299bf1c67d46ce2c/Diet-of-the-loggerhead-sea-turtles-in-the-Gulf-of-Gabes-Southern-Tunisia-Central-Mediterranean-Sea.pdf.

Lazar, B., Zavodnik, D., Grbac, I., & Tvrtkovic, N. (2002). Diet composition of the loggerhead sea turtle Caretta caretta in the northern Adriatic Sea: a preliminary study. Mosier A., Folley A., Brost, B. (Comps.). Proceedings of the 20th Annual Symposium on Sea Turtle Biology and Conservation, Orlando, Florida. NOAA technical memorandum NMFS-SEFSC-477.

Margaritoulis, D., Lourenço, G., Riggall, T.E., & Rees, A.F. (2022). Thirty-Eight Years of Loggerhead Turtle Nesting in Laganas Bay, Zakynthos, Greece: A Review. Chelonian Conservation and Biology 21(2), 143–157. https://doi.org/10.2744/CCB-1531.1.

Mariani, G., Bellucci, F., Cocumelli, C., Raso, C., Hochscheid, S., Roncari, C., Nerone, E., Recchi, S., Di Giacinto, F., Olivieri, V., Pulsoni, S., Matiddi, M., Silvestri, C., Ferri, N., & Di Renzo, L. (2023). Dietary Preferences of Loggerhead Sea Turtles (Caretta caretta) in Two Mediterranean Feeding Grounds: Does Prey Selection Change with Habitat Use throughout Their Life Cycle? Animals 13(4), 654. http://dx.doi.org/10.3390/ani13040654.

Mulochau, T., Jean, C., Gogendeau, P., & Ciccione, S. (2021). Green sea turtle, Chelonia mydas, feeding on Synapta maculata (Holothuroidea: Synaptidae) on a seagrass bed (Syringodium isoetifolium) at Reunion Island, western Indian Ocean. SPC Beche-de-mer Information Bulletin (41), 37–39.

Pajuelo, M., Bjorndal, K.A., Arendt, M.D., Foley, A.M., Schroeder, B.A., Witherington, B.E., & Bolten, A.B. (2016). Long-term resource use and foraging specialization in male loggerhead turtles. Marine Biology 163(235). https://doi.org/10.1007/s00227-016-3013-9.

Palmer, J.L., Beton, D., Çiçek, B.A., Davey, S., Duncan, E.M., Fuller, W.J., Godley, B.J., Haywood, J.C., Hüseyinoğlu, M.F., Omeyer, L.C.M., Schneider, M.J., Snape, R.T.A, & Broderick, A.C. (2021). Dietary analysis of two sympatric marine turtle species in the eastern Mediterranean. Marine Biology 168(6), 94. http://dx.doi.org/10.1007/s00227-021-03895-y.

Papafitsoros, K. (2015). In-water behaviour of the loggerhead sea turtle (Caretta caretta) under the presence of humans (Homo sapiens) in a major Mediterranean nesting site. Proceedings of the 35th annual symposium on sea turtle biology and conservation, Dalaman, Turkey.

Papafitsoros, K. (2022a). Social media mining and photo-identification detects the shift of longterm seasonal foraging habitat for a juvenile loggerhead sea turtle. MedTurtle Bulletin 1. https://doi.org/10.1101/2022.02.28.482324.

Papafitsoros, K. (2022b). Underreported in-water behaviours of the loggerhead sea turtle: Getting buried in the sand. MedTurtle Bulletin 2. https://doi.org/10.1101/2022.08.26. 505133.

Papafitsoros, K., Adam, L., & Schofield, G. (2023). A social media-based framework for quantifying temporal changes to wildlife viewing intensity. Ecological Modelling 476, 110223. https://doi.org/10.1016/j.ecolmodel.2022.110223.

Papafitsoros, K., Panagopoulou, A., & Schofield, G. (2021). Social media reveals consistently disproportionate tourism pressure on a threatened marine vertebrate. Animal Conservation 24(4), 568–579. https://doi.org/10.1111/acv.12656.

Papafitsoros, K. & Schofield, G. (2016). Focal photograph surveys: Foraging resident male interactions and female interactions at fish-cleaning stations. Proceedings of the 36th Annual Symposium on Sea Turtle Biology and Conservation, Lima, Peru. NOAA Tech Memo NMFS-SEFSC-734.

Rogers, A., Caal, W., Hamel, J.-F., & Mercier, A. (2020). Loggerhead sea turtle Caretta caretta found preying on a sea cucumber on a reef in Belize. SPC Beche-de-mer Information Bulletin (40), 17–19.

Schofield, G., Katselidis, K.A., Lilley, M.K.S., Reina, R.D., & Hays, G.C. (2017). Detecting elusive aspects of wildlife ecology using drones: New insights on the mating dynamics and operational sex ratios of sea turtles. Functional Ecology 31(12), 2310–2319. https://doi.org/10.1111/1365-2435.12930.

Schofield, G., Klaassen, M., Papafitsoros, K., Lilley, M., Katselidis, K.A., & Hays, G.C. (2020). Long-term photo-id and satellite tracking reveal sex-biased survival linked to movements in an endangered species. Ecology 101 (7), e03027. https://doi.org/10.1002/ecy.3027.

Schofield, G., Papafitsoros, K., Chapman, C., Shah, A., Westover, L., Dickson, L.C.D., & Katselidis, K.A. (2022). More aggressive sea turtles win fights over foraging resources independent of body size and years of presence. Animal Behaviour 190, 209–219. https://doi.org/10.1016/j.anbehav.2022.05.006.

Vander Zanden, H.B., Bjorndal, K.A., & Bolten, A.B. (2013). Temporal consistency and individual specialization in resource use by green turtles in successive life stages. Oecologia 173, 767–777. https://doi.org/10.1007/s00442-013-2655-2.

Vander Zanden, H.B., Bjorndal, K.A., Reich, K.J., & Bolten, A.B. (2010). Individual specialists in a generalist population: results from a long-term stable isotope series. Biology Letters 6(5), 711–714. http://dx.doi.org/10.1098/rsbl.2010.0124.

Vicente, V.P. & Carballeira, N. (1991). Studies on the feeding ecology of the Hawksbill Turtle Eretmochelys imbricata in Puerto Rico. Salmon M. and Wyneken J. (Comps.). Proceedings of the 11th Annual Workshop on Sea Turtle Biology and Conservation, Jekyll Island, Georgia. NOAA technical memorandum NMFS-SEFSC302.

